# Molecular Docking Analysis Of Some Phytochemicals On Two SARS-CoV-2 Targets: Potential Lead Compounds Against Two Target Sites of SARS-CoV-2 Obtained from Plants

**DOI:** 10.1101/2020.03.31.017657

**Authors:** Amaka Ubani, Francis Agwom, Oluwatoyin RuthMorenikeji, Shehu Nathan, Pam Luka, Arinze Umera, Usal Umar, Simeon Omale, Nnaemeka Emmanuel Nnadi, John Chineye Aguiyi

## Abstract

COV spike (S) glycoprotein and M^pro^ are two key targets that have been identified for vaccines and drug development against the COVID-19 disease. Virtual screening of some compounds of plants origin that have shown antiviral activities were carried out on the two targets, 6lu7 and 6vsb by docking with the PyRx software. The binding affinities were compared with other compounds and drugs already identified as potential ligands for 6lu7 and 6vsb as well as Chloroquine and hydroxychloroquine. The docked compounds with best binding affinities were also filtered for drug likeness using the SwissADME and PROTOX platforms on the basis of Physicochemical properties and toxicity respectively. The docking results revealed that scopodulcic acid and dammarenolic acid had the best binding affinity on the s-glycoprotein and M^pro^ protein targets respectively. Silybinin also demonstrated a good binding affinity to both protein targets making it a potential candidate for further evaluation as repurposed candidate for SARS COV2 with likelihood of having a multitarget activity.

## 1. Introduction

Severe acute respiratory syndrome coronavirus 2 (SARS-CoV-2), family *Coronaviridae*, genus *Betacoronavirus*, is spreading widely in China, causing coronavirus disease 2019 (COVID-19)(1). Since 2003, three Coronaviruses have been associated with pneumonia, the first was severe acute respiratory syndrome coronavirus (SARS-CoV)(2) which affected 8,098 people causing 774 deaths between 2002 and 2003(3), the second was Middle-East respiratory syndrome coronavirus (MERS-CoV)(4) which affected 27 countries and infecting a total of 2,494 individuals and claiming 858 lives(4). SARS-CoV-2 is a human pathogen which has been declared a global pandemic by the World Health Organisation (5). SARS-CoV and SARS-CoV-2 are closely related and originated in bats, who most likely serve as reservoir host for these two viruses (4). To date, no therapeutics or vaccines are approved against any human-infecting coronaviruses(4).

The entry into the host cell by the Coronaviruses is usually mediated by spike (S) glycoprotein (4). This glycoprotein interacts with the angiotensin-converting enzyme 2 (ACE2) enabling the virus penetration into the host. The main protease (M^pro^ also known as 3CL^pro^) is one of the best characterized drug targets among coronaviruses (6). The protease enzyme is essential for processing the polyproteins that are translated from the viral RNA(7). For this study, these two drug targets were selected for SARS-CoV-2 using plant based compounds screened against them.

Therefore, potent inhibitors of these two targets will be able to interfere with the SARS COV-2 replication process and thus serves as potential drugs for the management of the COVID-19. Hence, this work is aimed at identifying other potential lead compounds of plant origin that can serve as candidates for testing against the SARS COV2 virus.

## 2. Results

A library of 22 compounds of plant origin known to have antiviral activity were obtained from Pubchem data base. Though the compounds are chemically diverse they consist of largely flavonoids and terpenes. Some compounds from the citrus family made were found among the library, though they could not make it among the top six selected compounds for each target demonstrated some good binding affinities.

Most of the compounds has showed similar binding affinities to the selected protein targets (6lu7 and 6vsb) compared to the training sets of known ligands to the selected targets. (See table 1)

**Table 1.**
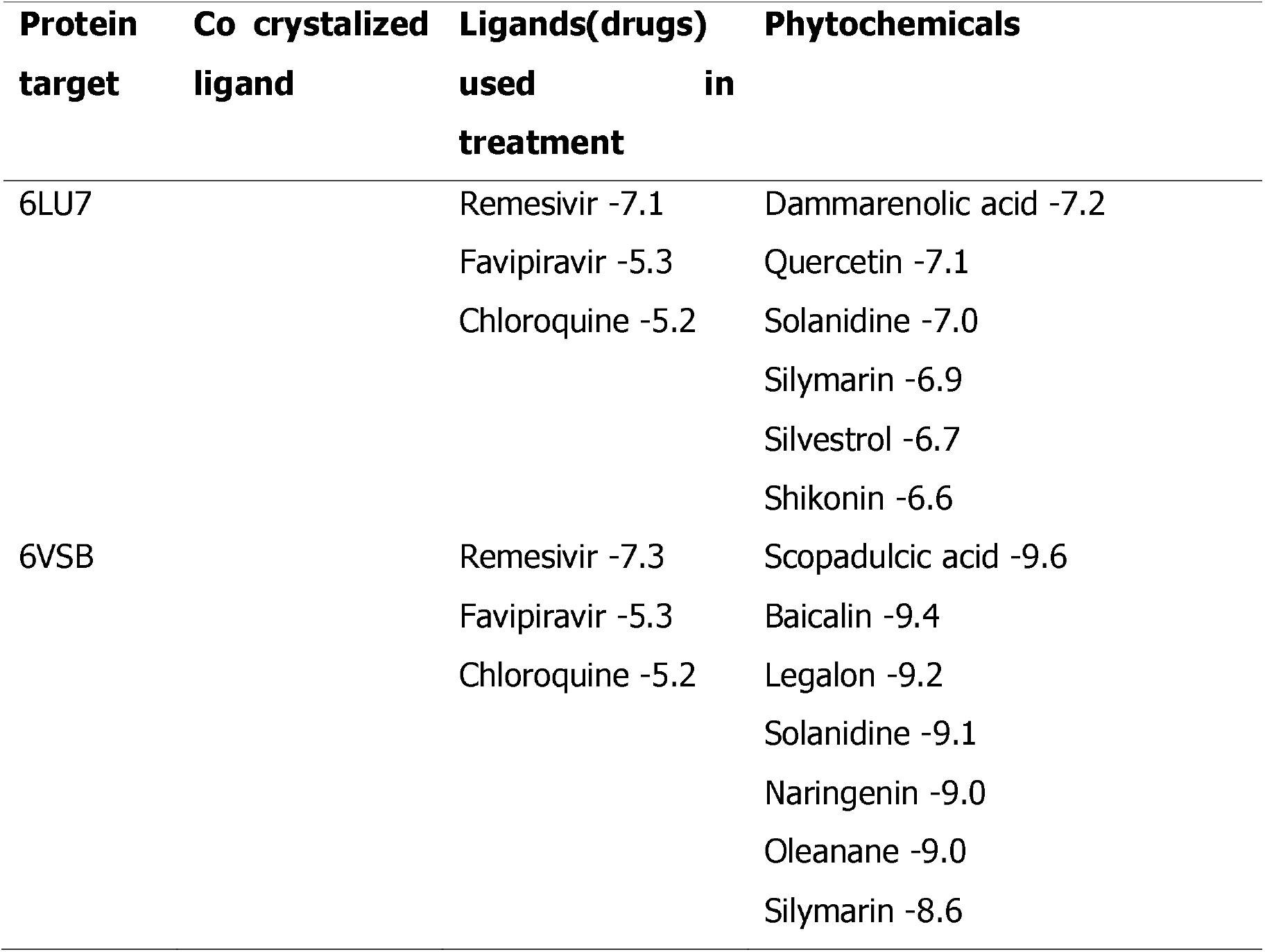
Comparison of Binding affinities of library to some known ligands and the co-crystalised ligand

However, the top six compounds with most favourable binding affinity were selected for each of the targets.

The outcome of the binding affinities of the selected compounds on the 6lu7 and 6vbs targets are presented in **Table 2 and Table 3** respectively.

**TABLE 2.**
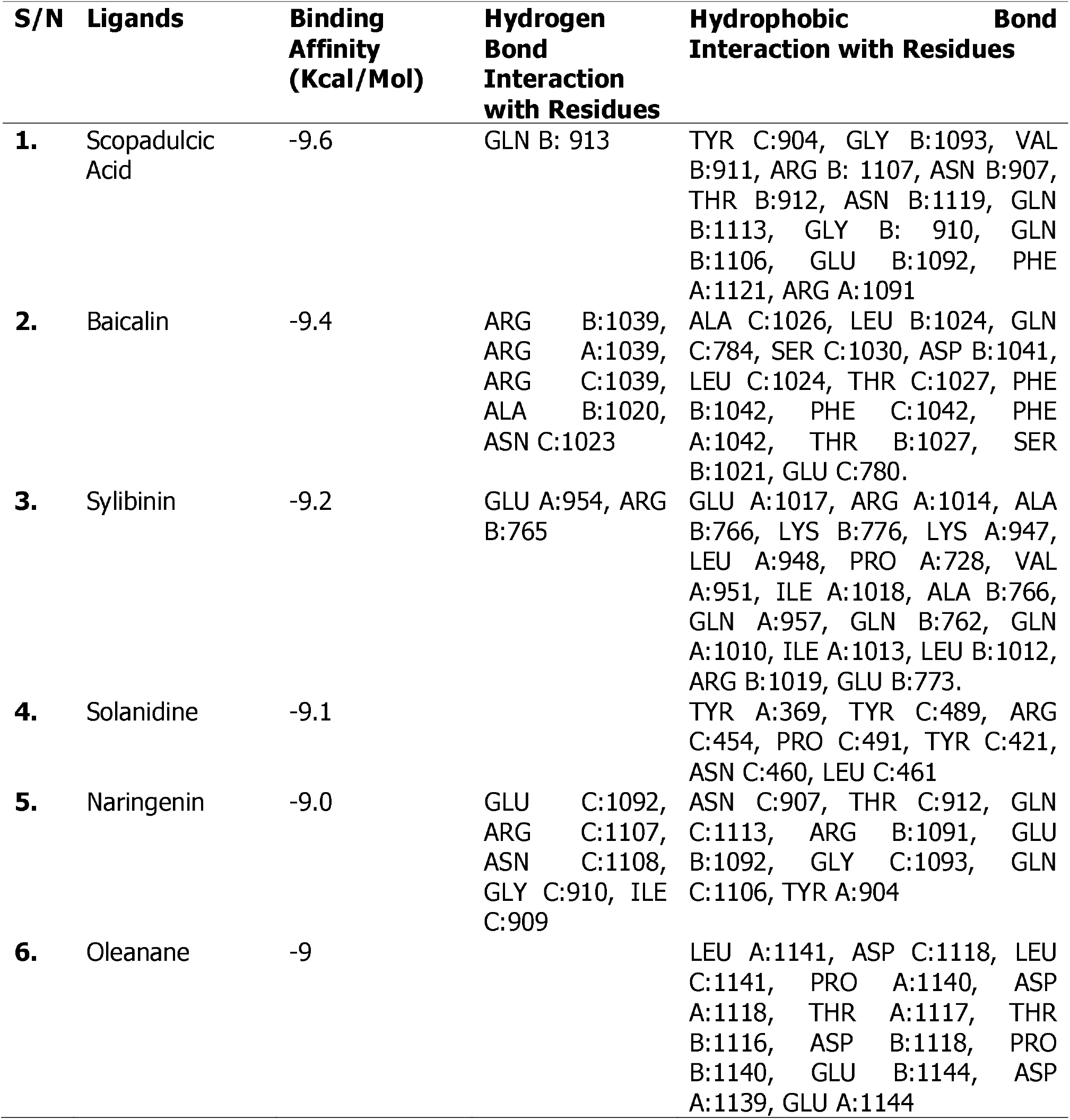
binding affinities of the compounds on the 6vsb and their Interaction with the binding site

**TABLE 3.**
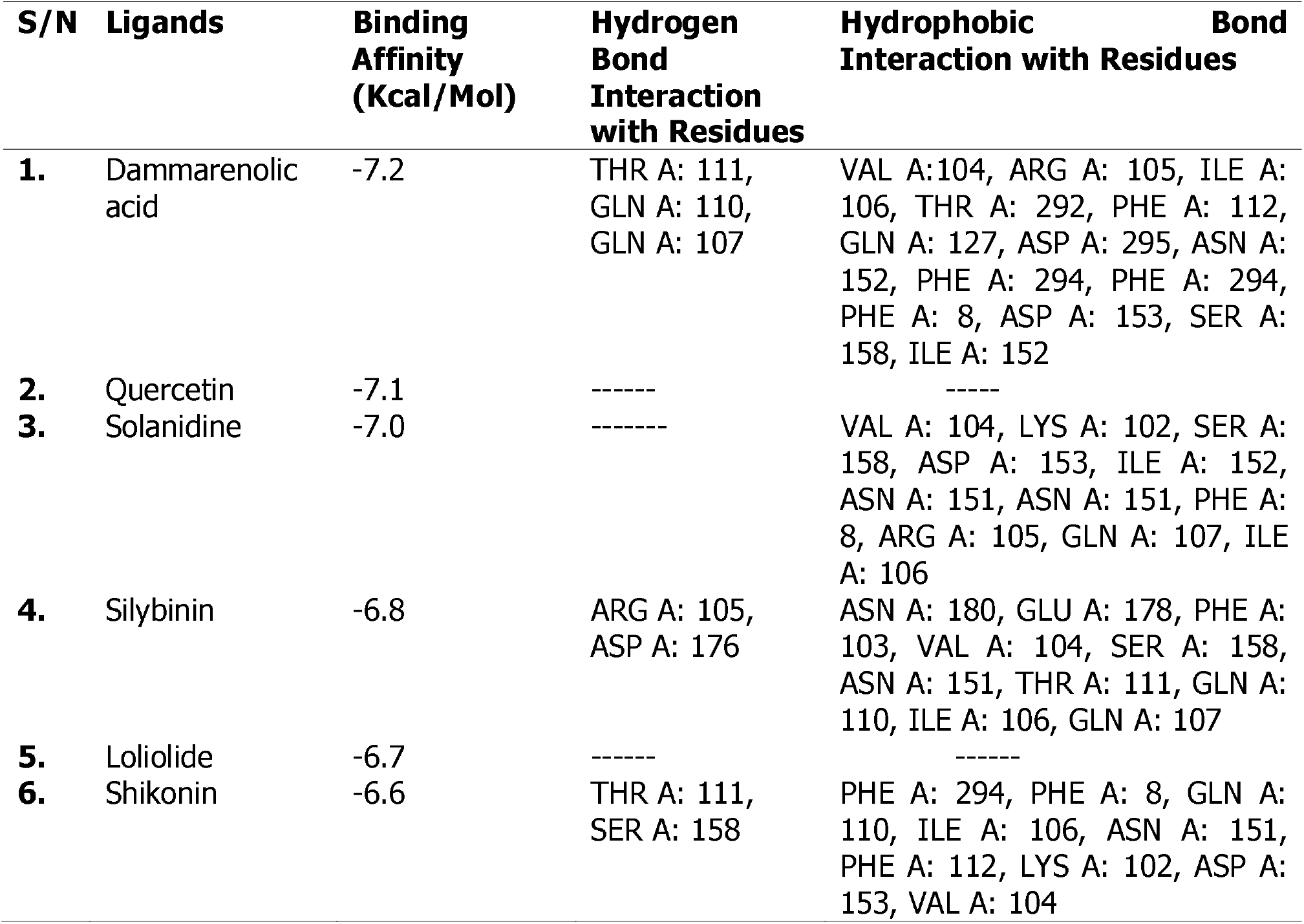
binding affinities of the compounds on the 6LU7 and their Interaction with the binding site

The binding affinities of the top six compounds on the 6vsb target are comparable to each other that is they all lie within a close range of 9 to 9.6 kcal/mol indicating that they might likely have equal or comparable potential as lead compounds for the 6vsb spike glycoprotein.

One of the compounds sylibinin (8)is an FDA approved drug, which showed up as active on both MPro and spike glycoprotein will make a good candidate of repurposing. Finding Quercetin as a potential inhibitor of the Mpro Protein (6flu7) of the SARS COV2 corresponds with an earlier report(9)

Looking at the the cLog P of the compounds, there was no correlation observed between the lipophilicity and the interaction with the receptors. However, for the compounds acting on 6lu7 (S/N 3,4,7,8,9 and 10 in Table 4), interaction with the receptor is correlated with low lipophilicity with the exception of solanidine and Dammarenolic acid that are have high cLogP values. Though both compounds also use their polar functional groups in interacting with the receptor. Bacailin and Naringenin showed good hydrogen bond interaction with the 6vsb receptor due to their polarity.

**Table 4.**
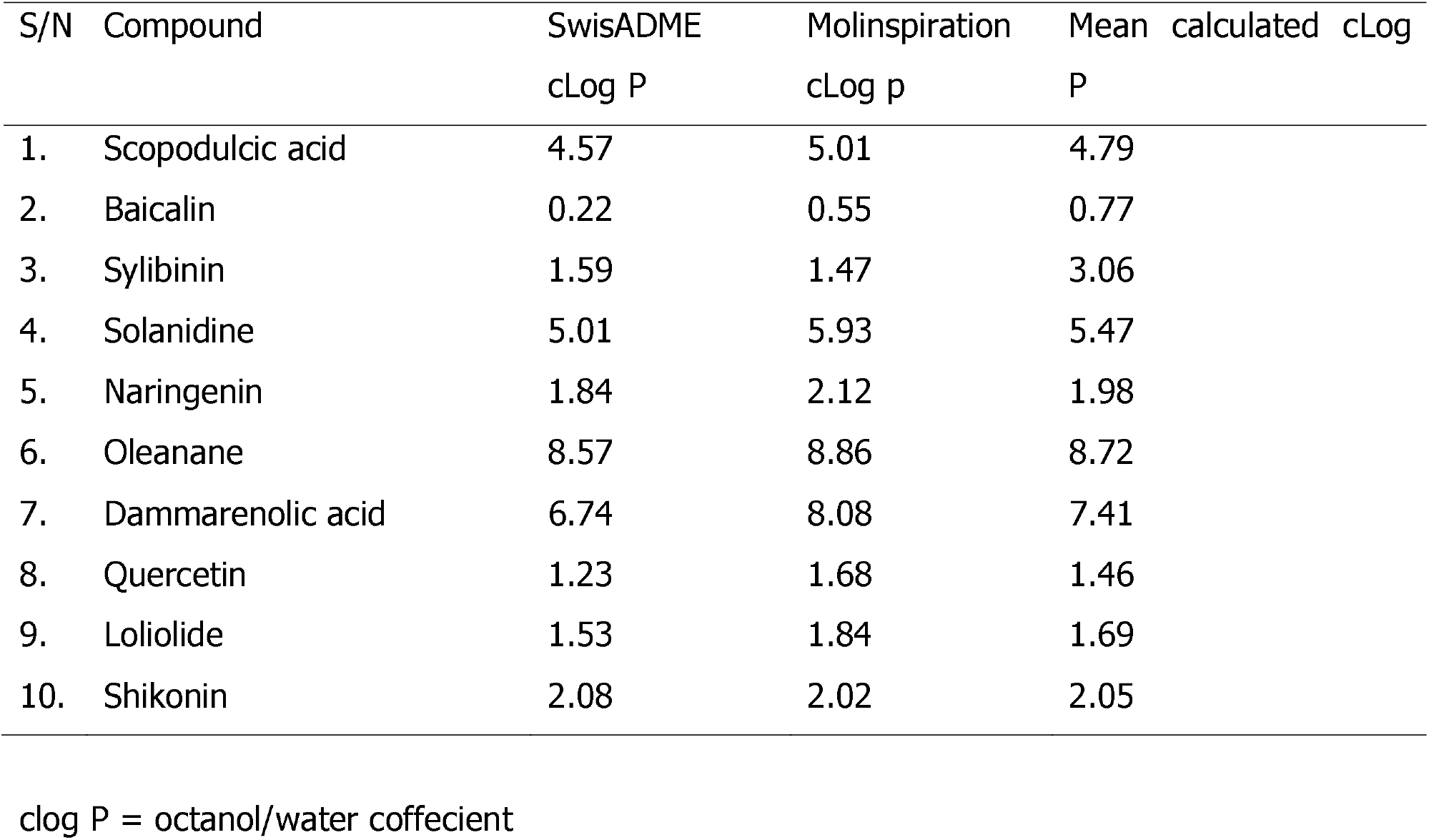
Comparison of the calculated cLog P values for the selected compounds

Filtering the compounds for drug likeness on the basis of Linpinski’s and/or Veber’s rule showed that all the compounds have drug like properties except baicalin which failed the two filtering scales applied (Table 5). This implies that baicalin is not worth considering further without any structural modification.

**Table 5.** Drug likeness.

| S/N | Compound | Mol.<br>Wt<sup>a</sup>(g/mol) | TPSA<sup>b</sup> | HBA<sup>c</sup> | HBD<sup>d</sup> | RB<sup>e</sup> | cLogP<sup>f</sup> | Lipinski<br>filter | Veber<br>filter |
| --- | --- | --- | --- | --- | --- | --- | --- | --- | --- |
| 1. | Scopodulcic<br>acid | 438.56 | 80.67 | 5 | 1 | 4 | 4.79 | + | + |
| 2. | Baicalin | 446.36 | 187.12 | 11 | 6 | 4 | 0.77 | - | - |
| 3. | Sylibinin | 482.44 | 155.14 | 10 | 5 | 4 | 3.06 | + | - |
| 4. | Solanidine | 397.64 | 23.47 | 2 | 1 | 0 | 5.47 | + | - |
| 5. | Naringenin | 272.25 | 86.99 | 5 | 3 | 1 | 1.98 | + | + |
| 6. | Oleanane | 412.75 | 0.00 | 0 | 0 | 0 | 8.72 | + | + |
| 7. | Dammarenolic<br>acid | 458.72 | 57.53 | 3 | 2 | 1 | 7.41 | + | + |
| 8. | Quercetin | 302.24 | 131.36 | 7 | 5 | 1 | 1.46 | + | + |
| 9. | Loliolide | 196.24 | 46.53 | 3 | 1 | 0 | 1.69 | + | + |
| 10. | Shikonin | 288.3 | 94.83 | 5 | 3 | 3 | 2.05 | + | + |

<sup>a</sup>Mol. Wt.: Molecular weight
<sup>b</sup>TPSA: Total Polar Surface Area
<sup>c</sup>No. HBA: Number of hydrogen bond acceptors
<sup>d</sup>No. HBD: Number of hydrogen bond donors
<sup>e</sup>No. RB: Number of rotatable bonds

**Figure 1.**
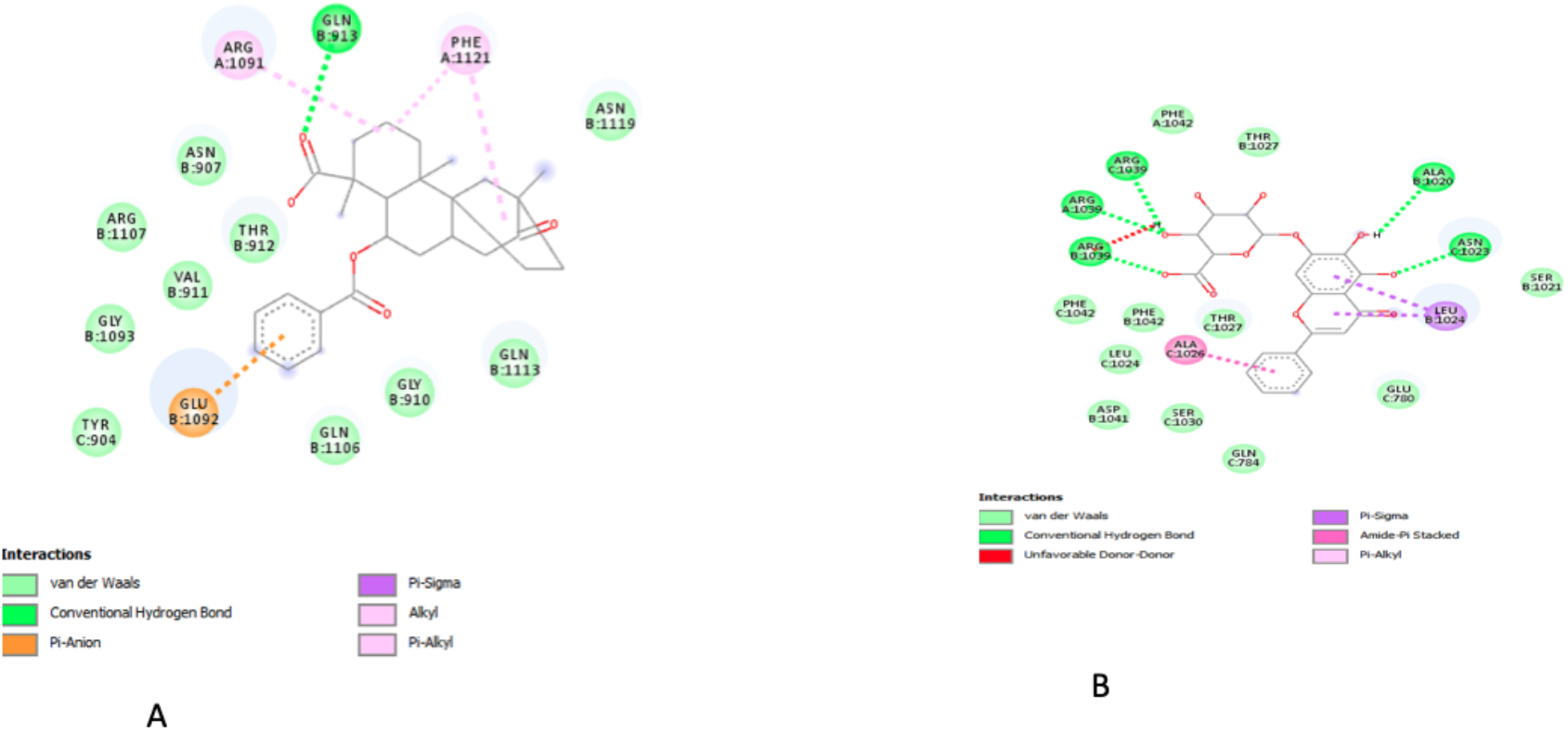
Best Binding pose and Interaction of (a) Scopodulcic acid and (b) Baicalin on the 6vsb protein.

**Table 6.**
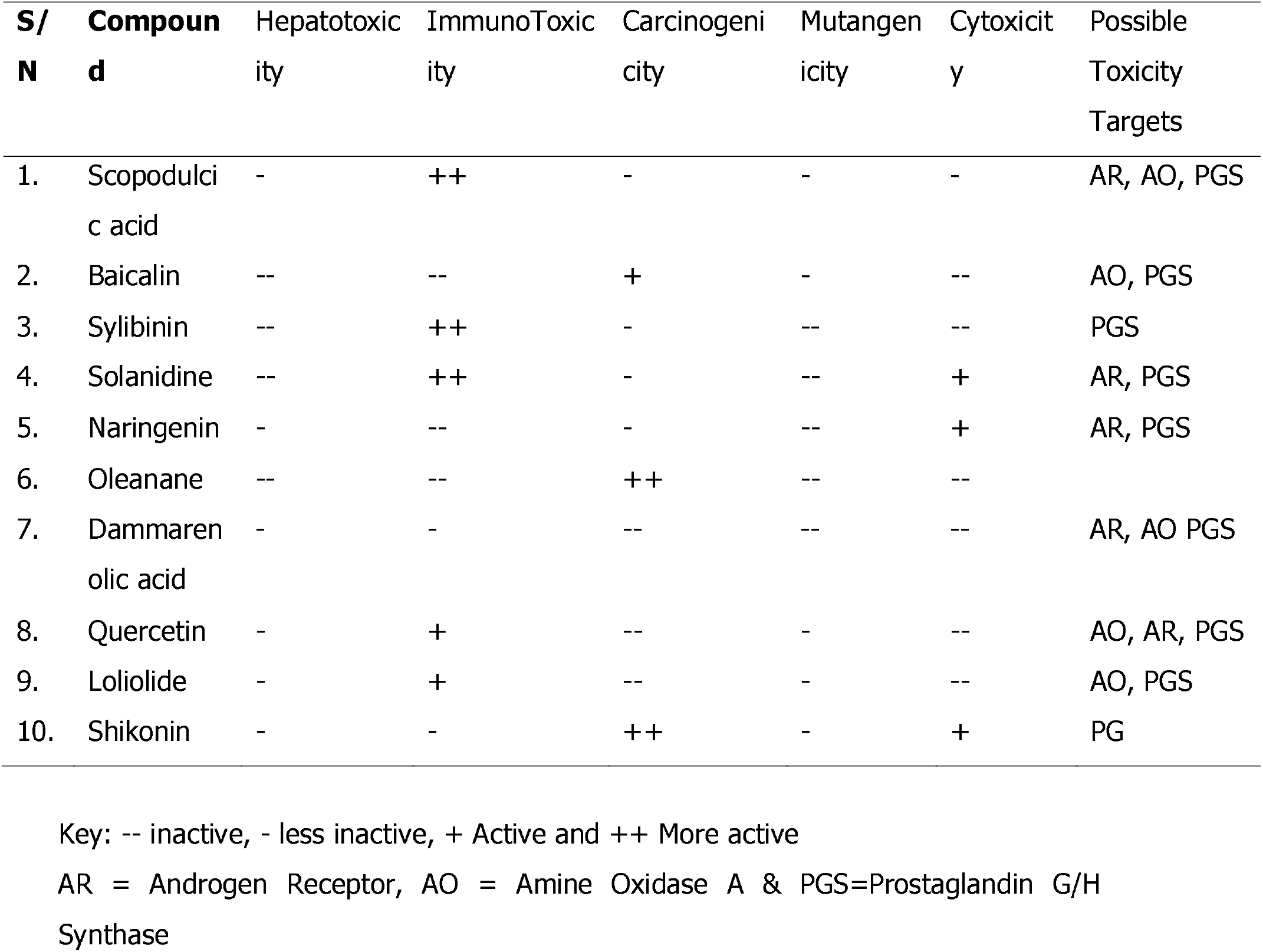
Predicted Toxicity Profile of the compounds using PROTOX II

The predicted toxicity profile of the selected compounds shows that all the compounds are likely to be relatively safe. Which makes them good potential candidates for anti-infectives because the chances of achieving selective toxicity is high. Baicalin is most likely the safest.

## Discussion

Two compounds among the top six selected for each target, solanidine and sylibinin were observed to have good binding affinity on both the 6vsb and the 6flu7. This make them potential multitarget acting inhibitors on the SARS-COV2. Solanidine is a Steroidal glycoalkaloids found in potatoes(10), although toxic to humans and animals, Solanidine has been reported to be effective against herpes viruses (HSV), herpes genitalis and herpes zoster(11) Its acivity against HSV is atttibuted to the presence of a sugar moiety(12). In silico method of drug screening using PROTOX II, showed that Solanidine is cytotoxic and immunotoxic. Prototox II is a cost and time conservative approach of testing and determining the toxicity of a compound to be considered a drug of choice(13). It incorporates molecular similarity, pharmacophores, fragment propensities and machine-learning models for the prediction of various toxicity endpoints; such as acute toxicity, hepatotoxicity, cytotoxicity, carcinogenicity, mutagenicity, immunotoxicity, adverse outcomes pathways (Tox21) and toxicity targets(13)

A safe drug must not be toxic to its host target. Based on the Protox II evaluation of Toxicity, Dammarenolic acid emerges as the compound of choice with the least toxicity. Dammarenolic acid have been reported as effective antiviral agents Dammarenolic acid potently inhibited the in vitro replication of other retroviruses, including Simian immunodeficiency virus and Murine leukemic virus in vector-based antiviral screening studies and has been proposed as a potential lead compound in the development of anti-retrovirals. (14) The compound is cytotoxic and demonstrate potential against respiratory syncytial virus(15). We therefore propose that the evaluation of Dammarenolic acid will hold the key to COVID19 drug considering its drugability and low toxicity.

This study proposes a potential re-purposing of silybinin for the management of COVID19 diseases. Silybinin(Silymarin) possesses potent antiviral activities against numerous viruses, particularly hepatitis C virus (HCV)(16, 17) It has been reported to have activities against a wide range of viral groups including flaviviruses (hepatitis C virus and dengue virus), togaviruses (Chikungunya virus and Mayaro virus), influenza virus, human immunodeficiency virus, and hepatitis B virus(16). Silymarin inhibits HCV in both *in vitro* and *in vivo* by inhibiting HCV entry, RNA synthesis, viral protein expression and infectious virus production; in addition it also acts by blocking of the virus cell-to-cell spread(18). As an FDA approved drug for the management of Hepatitis disease. In silico analysis of this drugs in this study has shown that it has acitivity against SAR COV 2 S-glycoprotein and proteas(M^pro^) targets making it a drug to be considered with multi-target ability in the management of this disease.

## 4. Materials and Methods

Plant Compounds with antiviral activities were mined from PubChem data base (https://pubchem.ncbi.nlm.nih.gov/). Two proteins including the main protease (6lu7) and the crystal structure of COVID-19 main protease(19) in complex with an inhibitor N3 and the Spike glycoprotein, N-ACETYL-D-GLUCOSAMINE (6vsb10.1126/science.abb2507) were downloaded from the protein database (PDB). The proteins were prepared using Discovery studio (version)(20) and a rigid docking scoring function was carried out using PyRx software(21). The results of the dock poses were visualized using Discovery Studio.

The Physicochemical properties and druggability of selected compounds were predicted using SwissADME(22) and Molinspiration(23) platforms and their predicted toxicity profile also compared using the PROTOX platform(24)

## 5. Conclusions

From the 22 phyto-compounds that were virtually screened, Scopodulcic acid and Dammarenolic acid showed the best binding energies with the Spike glycoprotein (6vsb) and the M^pro^ (6flu7) respectively. This makes them potential lead compounds for development into candidates against the SARS COV 2. Furthermore, the FDA approved drug silybinin (Legalon) with good binding affinity on the two targets can be evaluated further for possible repurposing against the SARS COV2 virus.

